# Bridging Ancestry Gaps in Genomic Risk Prediction with Tabular Foundation Models

**DOI:** 10.64898/2026.05.29.728877

**Authors:** Anirban Das, Yan Cui

## Abstract

**Motivation:** Models deployed for genomic prediction of diseases perform unevenly across populations, limiting clinical utility. Two factors drive this limitation: large imbalances in sample availability across ancestry groups and non-stationarity of genotype–phenotype effect sizes across the ancestry continuum. While tabular foundation models with in-context learning (ICL) have shown strong sample efficiency in other domains, their effectiveness for genotype-to-phenotype prediction and their robustness to ancestry-driven effect heterogeneity remain unclear.

**Results:** Using large, ancestrally diverse biobank data, we show that ICL-capable tabular foundation models reduce performance degradation in under-sampled ancestry groups compared to conventional supervised approaches. However, we find that prevailing models trained on existing synthetic tabular tasks fail when allele effect sizes vary across ancestry space. Treating genetic ancestry as a continuous variable, we introduce an instruction-tuning framework that exposes models to synthetic tasks with ancestry-dependent non-stationary effects. Instruction-tuned models achieve improved and more stable predictive performance across the genetic ancestry continuum, including for individuals distant from in-context exemplars in ancestry space.

**Availability and Implementation:** All code for instruction-tuning models, synthetic task generation, data wrangling, and model evaluation, is publicly available at https://github.com/ai4pm/Bridging-Ancestry-Gaps-in-Genomic-Risk-Prediction-with-Tabular-Foundation-Models. The final instruction-tuned model (ICL-NS-G2P-proto) is also released in this repository. Detailed documentation is provided, including environment setup instructions and guidelines for running various parts. The instruction-tuning task datasets are available at https://zenodo.org/records/18309187.

**Supplementary Information:** Supplementary data are available online.

## Introduction

Genomic prediction of diseases is a growing area of computational genomics with applications in disease screening and precision medicine. Despite steady gains in genome-wide association studies (GWAS) and polygenic risk score (PRS) methodology, predictive performance remains uneven across populations, limiting clinical utility and raising concerns about equity (Kullo, 2025). These limitations stem from structural properties of current genomic datasets and modeling assumptions that do not hold across diverse populations.

A primary barrier to equitable genomic prediction is the imbalance in sample availability across ancestry groups in biobank resources (Sirugo et al., 2019; Gurdasani et al., 2019; Mills and Rahal, 2020; Guerrero et al., 2018; Bien et al., 2019). As a result, supervised prediction models trained on these data exhibit reduced accuracy and increased uncertainty when applied to underrepresented groups, even for diseases with substantial heritable components (Duncan et al., 2019; Privé et al., 2022; Wang et al., 2020). Although recent multi-ancestry efforts have improved discovery and fine-mapping (Shrine et al., 2023; Cui, 2025; Tcheandjieu et al., 2022; Henry et al., 2025; Zhou et al., 2022), they do not by themselves resolve the sample-efficiency requirements of downstream prediction models.

A second challenge is that genotype–phenotype relationships are not stationary across populations. Empirical studies show that allele effect sizes and genetic architectures can vary across ancestry backgrounds and cohorts (Tcheandjieu et al., 2022; Shrine et al., 2023; Henry et al., 2025; Wang et al., 2023). These shifts violate assumptions common to many PRS and machine learning approaches—that a single set of effect estimates transfers unchanged between populations. As biobank studies expand to include broader and more diverse cohorts, this non-stationarity becomes increasingly consequential (Kachuri et al., 2023; Kullo, 2025; Gao et al., 2023).

### From supervised learning to ICL capability

Most genomic risk models are trained using conventional supervised learning, in which model parameters are optimized for a fixed dataset and task. When applied to new data such models require retraining from scratch. This paradigm is limiting (Wang et al., 2020) in genomic settings, where labeled data availability varies sharply across ancestry groups.

In-context learning (ICL) is a capability that has emerged in large foundation models, most prominently in language models, whereby a pretrained model adapts its predictions at inference time by conditioning on a small number of labeled examples provided as part of the input, without updating its parameters (Wei et al., 2022). Rather than learning a single task-specific mapping, an ICL-capable model is trained to infer task structure from in-context exemplars.

This capability can be acquired through instruction tuning, in which a foundation model is trained on a large collection of tasks. Each task is presented as a small set of labeled exemplars followed by unlabeled examples to be predicted, and the model’s parameters are updated across tasks to improve its ability to solve new tasks from few examples (Wei et al., 2022; Bai et al., 2023). Through exposure to many such tasks, the model learns a general strategy for learning from context.

Recent breakthroughs have led to the emergence of tabular foundation models capable of in-context learning (Hollmann et al., 2025). They have been shown to maintain strong predictive performance on small tabular datasets. These properties are particularly promising for genomic risk prediction in under-sampled ancestry groups. Two critical gaps remain: it is unknown whether the sample-efficiency advantages of ICL extend to genotype-to-phenotype prediction, and existing tabular foundation models are not designed to handle spatial non-stationarity in data generation processes.

### Modeling ancestry as a continuum

Human genetic variation is continuous and structured by admixture and fine-scale population history, yet many prediction pipelines rely on coarse ancestry labels (Peterson et al., 2019). Recent work demonstrates that PRS accuracy often decays smoothly with genetic distance from the training set, even within broadly defined ancestry groups (Ding et al., 2023). Treating ancestry as a continuum provides a principled way to quantify this decay, evaluate robustness as ancestry dispersion increases, and avoid artifacts introduced by discrete group boundaries (Ding et al., 2023; Kachuri et al., 2023). This perspective is particularly important when studying non-stationarity, as locally smooth genetic effects may appear stable in narrow ancestry regions but vary substantially across broader spans of ancestry space. Transfer learning (TL) is a common strategy for mitigating ancestry-related data imbalance by adapting a pretrained model to a target ancestry group or cohort (Gao and Cui, 2020; Gao et al., 2023; Gao and Cui, 2024; Sharma et al., 2026). However, TL is typically applied across a small number of discrete ancestry groups or cohorts, and does not extend to a continuous ancestry continuum.

### Overview and contributions

Motivated by these challenges, we study tabular foundation models for genotype-to-phenotype prediction in ancestrally diverse biobank data. We first evaluate whether the sample-efficiency advantages associated with in-context learning extend to genomic risk prediction under severe ancestry-related data imbalance. We then identify a critical limitation of existing tabular foundation models: sensitivity to ancestry-dependent non-stationarity in allele effect sizes. To address this gap, we align tabular foundation models to this setting using two complementary strategies: instruction tuning on synthetic tasks that explicitly encode non-stationary genetic effects, and representing ancestry as a continuous variable for both conditioning and evaluation. In the process, we introduce a novel family of synthetic genotype–phenotype prediction tasks designed to capture ancestry-dependent non-stationarity; such datasets are solely by themselves key resources for improving AI models (Das et al., 2025).

### In-context learning with tabular foundation models

Tabular foundation models such as TabPFN (Hollmann et al., 2025) can be deployed on new prediction tasks without any parameter updates, operating in a strict in-context learning (ICL) regime. The procedure is as follows. A new task is presented as a labeled training set (rows of feature vectors paired with known labels) together with an unlabeled test set whose labels are to be predicted. A conventional supervised model would be retrained from scratch on the training data before being applied to the test data. In contrast, TabPFN treats the labeled training samples as *in-context exemplars*: the training rows and the test rows are concatenated into a single input table, with the test labels masked. In one forward pass through the transformer, the model attends over the in-context exemplars to infer the task structure and produces a predictive distribution for every masked test label—without performing any gradient updates to its parameters. This strictly in-context procedure is used for all evaluations of both off-the-shelf TabPFN and ICL-NS-G2P-proto (the instruction-tuned variant derived from off-the-shelf TabPFN) reported in this study, unless stated otherwise.

## Results

### Benefits of in-context learning under ancestry-related sample imbalance

We evaluated whether the sample-efficiency advantages of in-context learning (ICL), well documented in language models (Wei et al., 2022; Brown et al., 2020), extend to genotype-to-phenotype prediction under ancestry-related data imbalance. The experiment centers on *phenotype–target ancestry–reference ancestry triplets*. For a given phenotype, the ancestry group with the largest training sample serves as the reference, while each remaining ancestry group is a target. Two quantities summarize each triplet. The Sample Availability Gap (SAG) measures the relative shortfall in training data between a target and reference ancestry: SAG = 1 − *n*_tgt_*/n*_ref_ . The Relative Performance Gap (RPG) measures how much a model’s predictive performance degrades for the target relative to the reference: RPG = (*M*_ref_ − *M*_tgt_) */M*_ref_, where *M* is the evaluation metric, either the ROC AUC or the lower bound of its 85% confidence interval (see Methods 3.2 for full details). For each triplet and metric, we compute the SAG and compare the RPG across three models (TabPFN, elastic net, and random forest), with the lowest RPG identifying the model least affected by sample imbalance.

Using data from the All of Us (AoU) Research Program, we analyzed 11 triplets spanning multiple disease categories. Across these comparisons, the ICL-capable TabPFN tended to exhibit smaller performance degradation, achieving the lowest RPG in 5 of 11 triplets for both metrics and never exhibiting the highest degradation (Fig. 1). To assess generalizability, we repeated the analysis using Phase III data from the NIH eMERGE Network (Lennon et al., 2022), evaluating six phenotypes: ACEI (ACE inhibitor–induced cough), CRF (cardiorespiratory fitness), DIV (diverticulosis), GERD (gastroesophageal reflux disease), HF (heart failure), and T2D (type 2 diabetes), with European (EUR) as reference and African (AFR) as target ancestry. The same qualitative pattern emerged: TabPFN showed the lowest RPG in 3/6 triplets (raw ROC AUC) and 5/6 triplets (lower bound CI), and the highest degradation in none (Fig. 2).

**Fig. 1.**
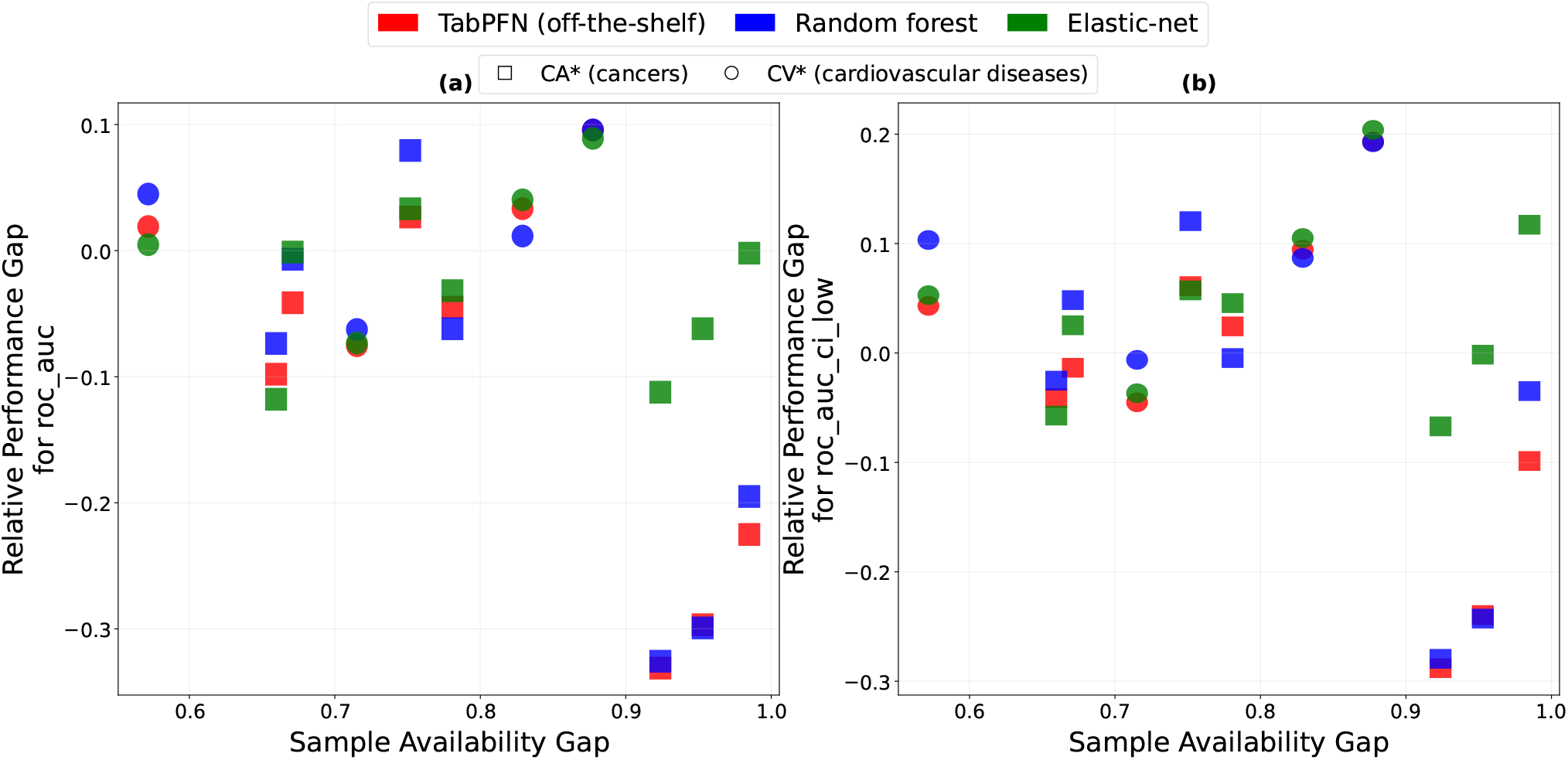
Relative Performance Gap (RPG, y-axis) versus Sample Availability Gap (SAG, x-axis) for three models across 11 phenotype–target–reference ancestry triplets. Triplets with cancer phenotypes are shown as rectangular markers and triplets with cardiovascular phenotypes as elliptical markers, data comes from AoU. Each triplet is evaluated using three models (TabPFN, elastic net, and random forest), which appear as three vertically aligned points sharing the same x-coordinate (different colors are used to represent the different models). Because these three points correspond to the same triplet, they share identical SAG; each point represents the RPG achieved by a different model. The lowest point on each vertical line indicates the model with the least performance degradation for that triplet, while the highest point indicates the model with the greatest degradation. (a) RPG computed using raw ROC AUC; (b) RPG computed using the lower bound of the 85% ROC AUC confidence interval. Triplet details are provided in the Supplementary Materials.

**Fig. 2.**
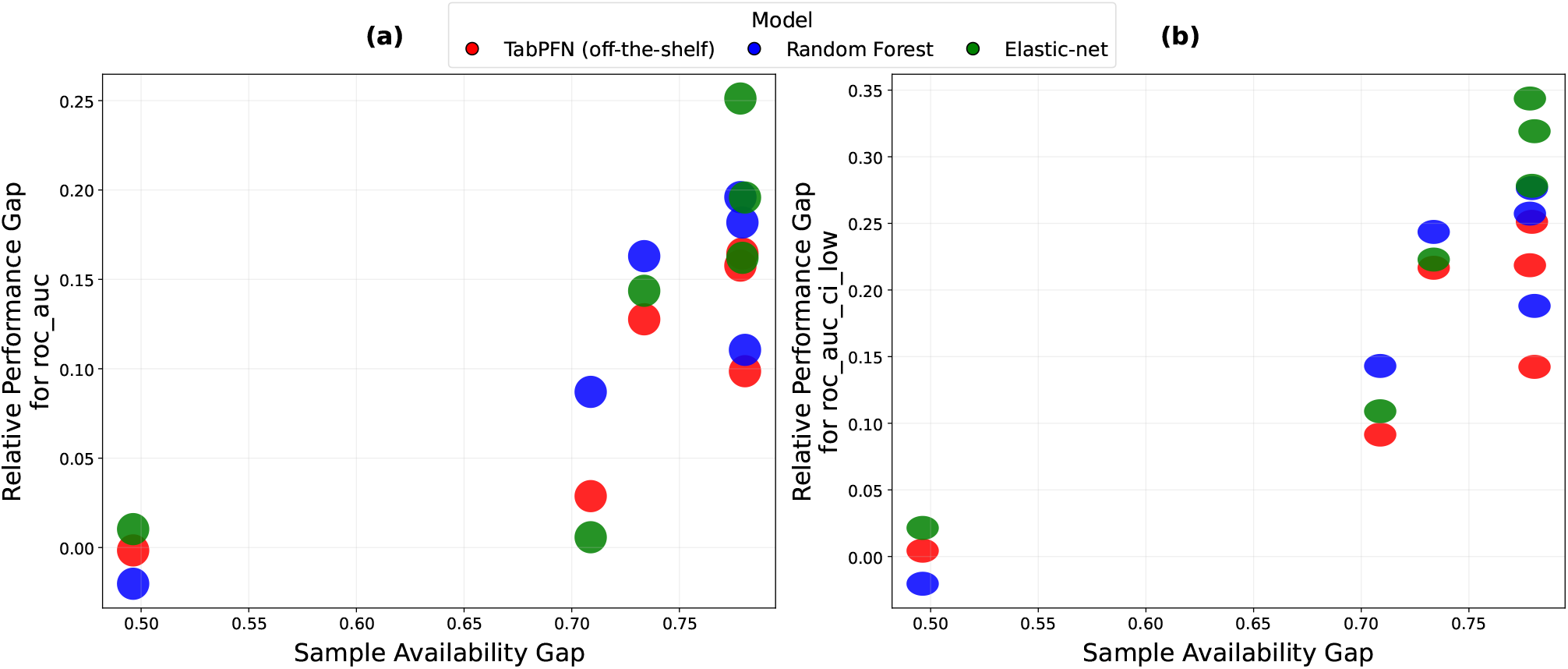
Same analysis as Fig. 1, applied to eMERGE data (see Fig. 1 caption for plot interpretation). (a) RPG computed using raw ROC AUC; (b) RPG computed using the lower bound of the 85% ROC AUC confidence interval.

These results demonstrate that tabular foundation models capable of in-context learning inherit a key advantage observed in other domains: improved robustness under limited labeled data. This property holds consistently across two independent biobank resources, multiple disease phenotypes, multiple ancestry contrasts, and multiple evaluation metrics, supporting the conclusion that ICL provides a mechanism for mitigating ancestry-related sample imbalance in genomic risk prediction.

### Identifying non-stationarity gaps in ICL-capable tabular foundation models

The results above indicate that ICL-capable tabular foundation models can mitigate ancestry-related performance degradation driven by sample imbalance. We next asked whether off-the-shelf tabular ICL-capable models remain effective if the data-generation process is non-stationary across ancestry space, i.e., genotype–phenotype effect sizes vary as a function of ancestry.

To isolate the impact of ancestry-dependent non-stationarity, we first evaluated model behavior in a controlled synthetic setting designed to mimic genotype–phenotype prediction across a continuous ancestry continuum. Each individual was assigned ancestry coordinates **r** ∈ ℝ^*A*^, and a vector of genotype dosages **x** = (*η*_1_, …, *η*_*N*_) ∈ ℝ^*N*^. Phenotype labels were generated according to a Bernoulli model with logistic link,

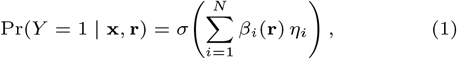

where *σ*(·) denotes the logistic function and the effect sizes *β*_*i*_(**r**) are unobserved functions of ancestry.

To explicitly induce non-stationarity, effect sizes for a subset of *N*_ctrl_ loci were modeled as Gaussian processes over ancestry space,

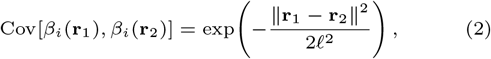

with length-scale parameter *ℓ*, controlling the smoothness of ancestry-dependent variation. Increasing *N*_ctrl_ introduces more loci with ancestry-varying effects, while smaller values of *ℓ*, induce sharper local heterogeneity (details in Supplementary).

In this setting, accurate prediction requires more than interpolation from in-context exemplars: the model must adapt as the target individual’s ancestry shifts relative to the exemplars. We find that off-the-shelf TabPFN fails to track these ancestry-dependent changes in effect size. Predictive performance deteriorates sharply as *N*_ctrl_ increases, with recovery only when *ℓ*, is sufficiently large that effect sizes are nearly constant across ancestry space (Fig. 3a). These results are expected as TabPFN is instruction-tuned on synthetic tasks that do not encode ancestry-dependent effect-size variation (Hollmann et al., 2025).

**Fig. 3.**
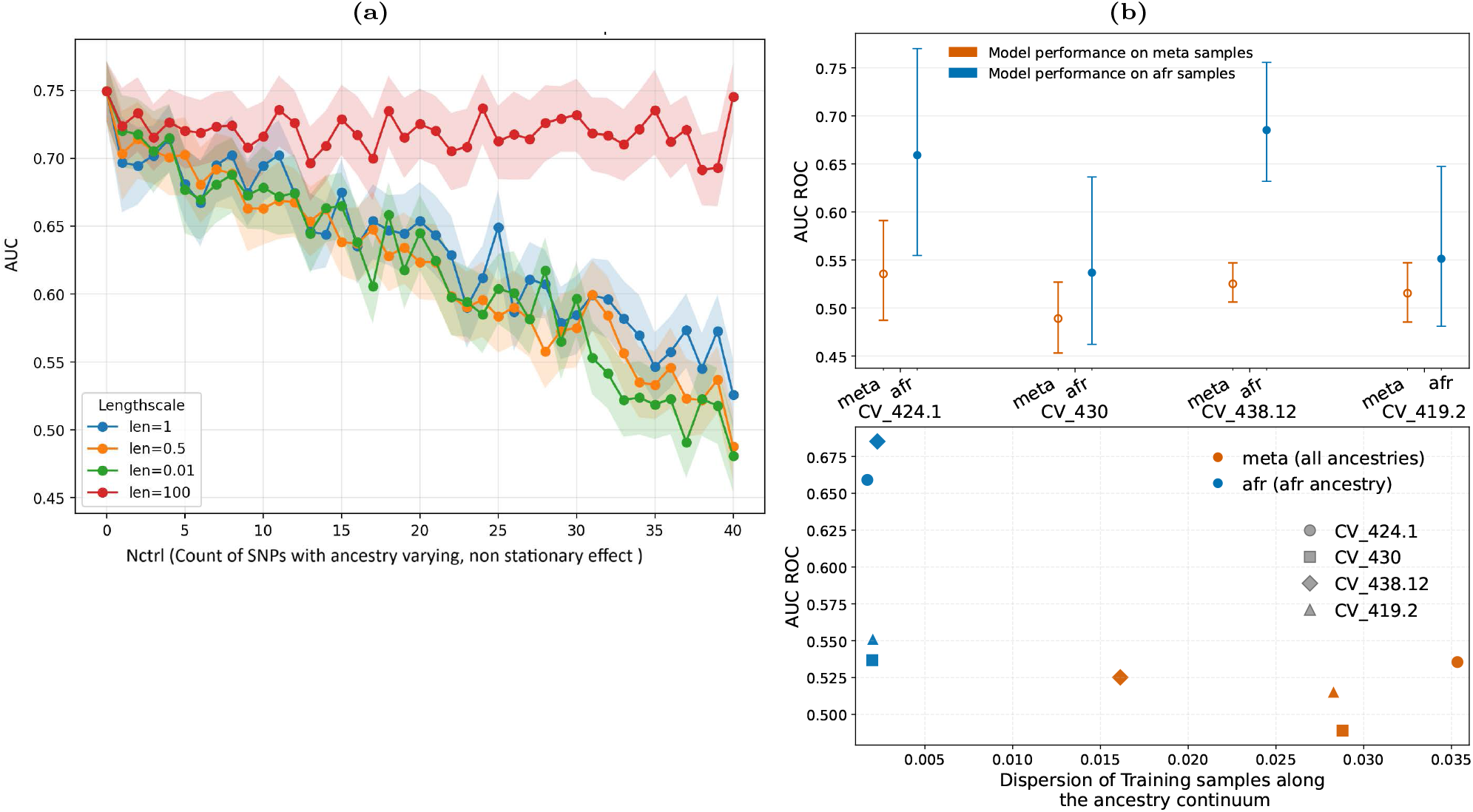
Non-stationarity gap in off-the-shelf TabPFN for genomic risk prediction. (a) Controlled evaluation with ancestry-dependent non-stationary effect sizes across a continuous ancestry coordinate. (b) Real-world biobank analyses (on AoU) showing systematic performance degradation across ancestry space when effect sizes vary with ancestry (top and bottom panels correspond to the same phenotype–ancestry settings).

We next evaluated whether the same limitation manifests in real-world biobank data, where ancestry-dependent non-stationarity arises from demographic history, linkage disequilibrium differences, and heterogeneous genetic architectures. Importantly, non-stationarity is expected to act as a limiting factor primarily when samples are drawn from a broad region of ancestry space; when ancestry dispersion is narrow, allele effect sizes are locally constant and prediction is less affected.

To test this, we selected four cardiovascular phenotypes, CV_430 (nontraumatic intracranial hemorrhage), CV_419.2 (presence of cardiac defibrillator), CV_438.12 (thoracic aneurysm), and CV_424.1 (left heart failure), and compared the performance of an off-the-shelf TabPFN model for each phenotype across two cohorts with distinct ancestry dispersion profiles. Individuals in the META (subjects of all ancestry groups combined) cohort were widely dispersed across ancestry space, whereas individuals in the AFR cohort formed a tightly clustered ancestry distribution Fig. 3b (bottom).

We observed that off-the-shelf TabPFN achieved substantially higher predictive performance in the AFR cohort than in the META cohort (Fig. 3b). This performance difference is consistent with the hypothesis that broader ancestry dispersion creates a setting in which ancestry-dependent non-stationarity of allele effect sizes—if present—would be expected to impact disease classification. Under this interpretation, the reduced performance of TabPFN in the ancestrally diverse META cohort reflects its inability to adapt predictions as genetic effect sizes vary across ancestry space. Taken together, these results indicate that off-the-shelf TabPFN fails to accommodate effect-size non-stationarity in real-world biobank data. Off-the-shelf tabular foundation models are not aligned to the non-stationary effect-size patterns that characterize genomic prediction across the ancestry continuum. This motivates an alignment step that exposes the model to ancestry-dependent non-stationarity during training, which we address next.

### Instruction tuning yields tabular foundation models that perform well across the ancestry continuum

To address the non-stationarity gap, we instruction-tuned the off-the-shelf TabPFN model on synthetic genotype–phenotype prediction tasks that explicitly encode ancestry-dependent variation in allele effect sizes, yielding an instruction-tuned prototype we refer to as ICL-NS-G2P-proto. Instruction tuning follows a standard in-context learning framework: the model is exposed during training to a large collection of tasks, each consisting of a small table of labeled in-context exemplars followed by unlabeled samples whose labels must be predicted. The task distribution used here differs from that of the original TabPFN in that it systematically incorporates non-stationary genotype–phenotype relationships across a continuous ancestry space (see Methods 3.4 for more details).

We evaluated ICL-NS-G2P-proto directly (see Methods 3.5) in an in-context learning regime, without fine-tuning on real data, focusing on meta-ancestry cohorts from the All of Us Research Program that were previously challenging for the off-the-shelf model. Across all four phenotypes, instruction tuning led to consistent improvements in predictive performance relative to the baseline TabPFN. These gains were observed across multiple metrics, including higher ROC AUC and lower cross-entropy loss, along with improved upper and lower confidence bounds (Fig. 4a). These improvements were achieved using a limited set of synthetic training tasks, indicating that exposing the model to ancestry-dependent non-stationarity during instruction tuning is sufficient to improve performance in ancestrally diverse cohorts.

**Fig. 4.**
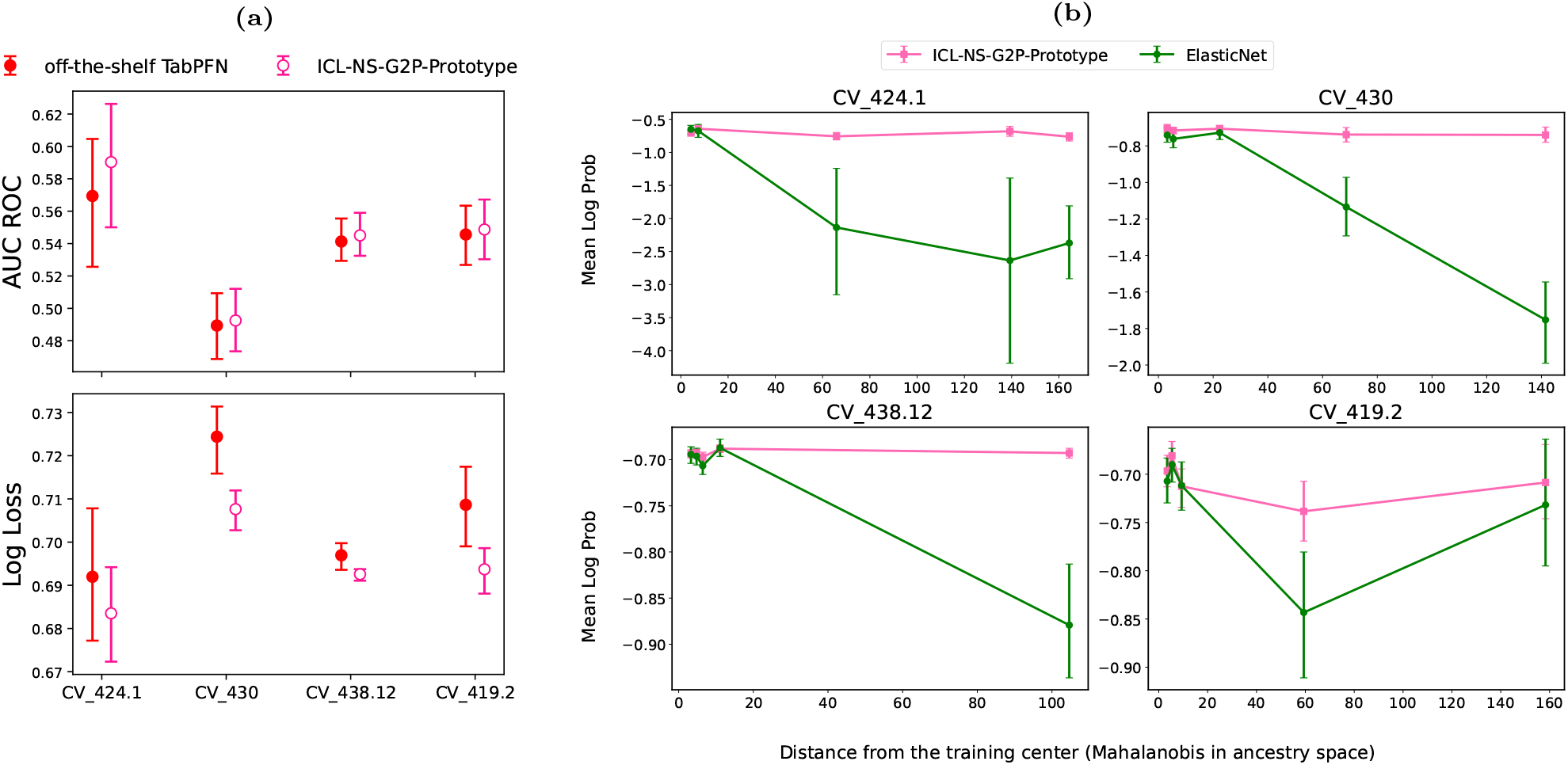
Instruction tuning yields tabular foundation models that perform well across the ancestry continuum. (a) Instruction-tuned ICL-NS-G2P-proto achieves higher ROC AUC and lower cross-entropy loss than off-the-shelf TabPFN across evaluated phenotypes in meta-ancestry cohorts (same AoU data as Fig 3). (b) Predictive performance as a function of ancestry distance from the training set. While elastic net performance deteriorates rapidly as distance increases, ICL-NS-G2P-proto maintains substantially more stable performance for samples far from in-context exemplars.

To further evaluate the ability of ICL-NS-G2P-proto to mitigate non-stationarity in ancestry-dependent genotype– phenotype relationships, we adopted an analysis framework introduced by Ding et al. (2023), which is applicable when ancestry is treated as coordinates in a continuous space. The underlying reasoning is as follows: a model that assumes stationarity will, during training, learn variant–phenotype relationships for individuals near the center of the training distribution. As test samples lie progressively further from this center in ancestry space, the learned genotype–phenotype relationships become increasingly inaccurate, because non-stationarity causes the true relationships to diverge from those learned in the training region. Consequently, models that cannot account for non-stationarity exhibit greater prediction error for subjects more distant from the training center. We constructed such a training–test split by restricting in-context exemplars/training-samples to individuals of EUR ancestry and evaluating predictions on test samples drawn from the mixed meta-ancestry cohort (see Methods 3.5). We quantified each test individual’s distance from the training distribution using the Mahalanobis distance between the individual’s ancestry coordinates and the center of the training set. We observed that the predictive performance of ICL-NS-G2P-proto deteriorated substantially less for samples increasingly distant from the training center compared to elastic net, across all four phenotypes examined (Fig. 4b). This demonstrates that ICL-NS-G2P-proto is robust to ancestral diversity in the test sample.

## Methods

### Preparing the All of Us dataset

We constructed genotype-to-phenotype prediction datasets from the All of Us (AoU) Research Program (Choi et al., 2024) within the AoU Researcher Workbench (Controlled Tier). The data-curation workflow and the primary AoU resources linked by our pipeline are summarized in the Supplementary Material.

For each phenotype, our pipeline produces a tabular dataset whose rows correspond to participants and whose columns include: (1) allele dosages for a phenotype-specific set of variants, (2) continuous ancestry coordinates derived from AoU-provided genetic principal components, (3) standard covariates such as age and sex, and (4) Disease labels.

#### All of Us All-by-All GWAS resources

A central component of the pipeline is the AoU-hosted All-by-All GWAS study conducted by the Broad Institute, in which genome-wide association analyses of AoU data was performed for 2,333 phenotypes across six ancestry groups, as well as in a meta-analysis combining all subjects. We focus on phenotypes in the phecodeX category for which at least 200 cases were observed in more than two ancestry groups. All analyses in the All-by-All study were performed using AoU version 7 data. We retrieve the 50 most significant associated variants for each disease–ancestry combination, do some post processing steps (Choi et al., 2020). These variants form the genetic feature set used to construct tabular datasets.

#### Managing data leakage and ancestry representation

Because All-by-All GWAS uses phenotype labels to identify associated variants, preventing leakage between feature discovery and evaluation is essential. We enforce strict subject-level separation by drawing training samples exclusively from AoU version 7 and evaluating on independent subjects who entered the program in AoU version 8. This ensures that no individual contributing to GWAS feature selection appears in the held-out evaluation set. To represent ancestry as a continuum rather than discrete labels, we use the coordinates provided by AoU as continuous ancestry coordinates (details in Supplementary Material). We implement a validation scheme to ensure the curated datasets and linkage steps produce reliable genotype-to-phenotype tables and that observed predictive performance is not driven by artifacts (details in Supplementary Material).

### Quantifying sample-imbalance effects via sample availability and relative performance gaps

To measure how predictive performance degrades in under-sampled ancestries, we compare each target ancestry group against a well-sampled reference ancestry for the same phenotype, and quantify degradation as a function of relative training-set size differences. This analysis is performed separately in All of Us (AoU) and in an independent biobank (eMERGE Phase III), using the same evaluation metrics and definitions across datasets.

#### AoU models and evaluation

TabPFN is evaluated in a strictly in-context learning (ICL) regime as described in Sec. 1.4: labeled AoU version 7 samples are provided as in-context exemplars, and the model predicts labels for held-out AoU version 8 individuals given only their features. In contrast, supervised baselines (elastic net and random forest) are trained from scratch on AoU version 7 and evaluated on AoU version 8. Model performance is summarized using ROC AUC, and uncertainty is quantified via bootstrap resampling of the test set; in addition to the point estimate, we report a conservative metric derived from the lower bound of an 85% confidence interval (using 100 bootstrap samples).

#### Sample Availability Gap (SAG) and Relative Performance Gap (RPG)

For each phenotype, we designate the ancestry group with the largest training sample size as the *reference* ancestry and treat all other ancestry groups as *target* ancestries. We define:

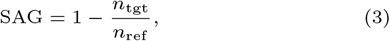

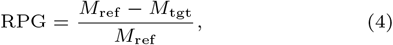

where *n*_ref_ and *n*_tgt_ are the reference and target training sample counts, respectively, and *M*_ref_ and *M*_tgt_ denote the chosen evaluation metric (e.g., ROC AUC or the lower bound of its 85% CI) computed on test sets restricted to the reference or target ancestry. Larger SAG indicates greater relative data scarcity, while larger RPG indicates greater relative performance degradation.

#### eMERGE dataset construction and phenotypes

We perform an evaluation in eMERGE Phase III using the same models and SAG/RPG definitions. Details of data wrangling are provided in the Supplementary Material. In Fig. 2 we report results for the six phenotypes: ACEI (ACE inhibitor–induced cough), CRF (cardiorespiratory fitness), DIV (diverticulosis), GERD (gastroesophageal reflux disease), HF (heart failure), and T2D (type 2 diabetes).

### Accounting for ancestry-specific heritability as a potential confounder

Beyond sample availability, predictive performance can vary across ancestry groups because the same phenotype may exhibit different SNP-based heritability when estimated from genomic features alone. Prior analyses within the AoU program document substantial ancestry-specific variation in disease heritability, raising the possibility that cross-ancestry performance differences could be driven in part by underlying differences in genetic signal rather than model behavior.

To mitigate this confounding factor, all model comparisons are performed *within* the same phenotype–target ancestry– reference ancestry triplets. We deliberately include phenotypes spanning a wide range of target-to-reference ancestry heritability ratios to assess robustness across heterogeneous genetic architectures. The phenotypes, ancestry groupings, sample sizes, and ancestry-specific heritability estimates are provided in the Supplementary Material. Ancestry-specific heritability estimates are not available for the eMERGE Phase III cohort.

### Isolating genetic non-stationarity along the ancestry continuum

For the analyses underlying Fig. 3 and Fig. 4, our objective is to isolate the impact of ancestry-dependent non-stationarity in genetic effects across the continuous ancestry space. Accordingly, we adopt a dataset construction strategy that minimizes the influence of non-genetic covariates and other potential confounders. For both the training dataset (AoU version 7 participants) and the test dataset (AoU version 8 participants), tabular representations exclude non-genetic covariates such as age and sex. Models are therefore restricted to learning from genetic features and continuous ancestry coordinates alone. To prevent confounding due to differences in covariate distributions, we enforce matching of case and control groups *within each dataset* with respect to the marginal distributions of age and sex. In addition, the number of cases and controls is matched within both the training and test sets.

### Instruction tuning for ancestry-aware genotype–phenotype prediction

#### Instruction-tuning overview

To develop ICL-NS-G2P-proto model, we adopt a standard instruction-tuning framework for in-context learning (ICL) (Wei et al., 2022), in which a base model is trained to generalize across families of tasks. We initialize from the publicly released TabPFN model (Hollmann et al., 2025) as the base tabular foundation model and retrain it on a large collection of newly generated synthetic tasks designed to capture ancestry-dependent non-stationary genotype–phenotype relationships.

During instruction tuning, a task is sampled uniformly from the family of synthetic genotype–phenotype tasks (see Sec. 3.4.2). For each sampled task, synthetic tabular data is generated according to the task’s data-generating process. The generated samples are then randomly split into in-context exemplars and test samples, with the proportion allocated to in-context exemplars varying uniformly between 70% and 90% for each task. The labels of the test samples are masked. The model processes the combined table using the in-context execution procedure described in Sec. 1.4: it attends over the labeled in-context exemplars to infer task structure and outputs predictions for the unlabeled entries. Training loss is computed using cross-entropy on these predicted labels. Model parameters are updated by accumulating loss across batches containing many such tables and performing gradient-based optimization. After one or more gradient updates on the current task, a new task is sampled from the task distribution, and training proceeds for a predetermined number of epochs. All model parameters are updated during instruction tuning.

#### Task generation via non-stationary genotype–phenotype models

The methodological novelty lies in the design of the task distribution. We could leverage the same class of synthetic genotype–phenotype models used to diagnose limitations of the base TabPFN model as generators of instruction-tuning tasks. Specifically, Gaussian process (GP) models are used to define ancestry-dependent, non-stationary allele effect sizes (Eq. 1 and Eq. 2). Each fixed configuration of GP parameters viz the length-scale parameter *ℓ*, and the number of loci with non-stationary effects *N*_ctrl_ —defines a distinct prediction task. For each task, labeled genotype–phenotype pairs can be generated, organized into learning tables, and used for instruction tuning. By varying GP parameters, we can generate a family of tasks. For each task, the resulting tabular data contains the following fields: ancestry coordinates of samples, genetic dosage features of samples, and, using Eqs. 1 and 2, a phenotype label.

Simple GP models do not fully capture several key characteristics of genotype–phenotype relationships in ancestrally diverse populations (Lee et al., 2025; Tcheandjieu et al., 2022). To better reflect these complexities, we instead employ hierarchical Gaussian process models to generate instruction-tuning tasks. In this formulation, individuals are sampled from a small number of clusters in ancestry space, representing coarse population structure. Allele frequencies and effect sizes are allowed to vary both between clusters and within clusters, with larger-scale variation modeled across clusters and finer-scale variation retained locally. Sample-density imbalances, such as oversampling from some ancestry clusters and undersampling from others, are explicitly incorporated. Further details of the hierarchical GP construction are provided in the Supplementary Material.

#### Obtaining ICL-NS-G2P-proto from off-the-shelf TabPFN

By varying hierarchical GP parameters, we construct a large and diverse family of genotype–phenotype prediction tasks spanning different regimes of ancestry-dependent non-stationarity, stratification, and sample imbalance. A subset of these tasks (190 in total) is used for instruction tuning of the base TabPFN model. To avoid information leakage during model development, a disjoint set of 30 tasks were generated using parameter configurations not seen during gradient updates. This set is reserved exclusively for hyperparameter selection, including learning rate, number of training epochs, and the number of gradient updates performed before switching tasks. The resulting instruction-tuned model is referred to as ICL-NS-G2P-proto.

### Evaluation of ICL-NS-G2P-proto within the biobank pipeline

We evaluate the instruction-tuned model (ICL-NS-G2P-proto) and the off-the-shelf TabPFN in an in-context learning (ICL) regime on AoU data constructed using the pipeline described in Sec. 3.1. In all evaluations below, neither tabular foundation model is fine-tuned on real training data; instead, labeled training samples are provided only as in-context exemplars and predictions are generated for held-out test samples.

For Fig. 4a, we fix the ancestry setting to the meta-ancestry cohort, where off-the-shelf TabPFN was observed to struggle due to non-stationarity in genotype–phenotype relationships. Using AoU version 7 participants as the training set and AoU version 8 participants as the test set, we construct phenotype-specific tabular datasets following the pipeline in Sec. 3.1. Predictive performance is summarized using multiple metrics, including area under the receiver operating characteristic curve (ROC AUC) and cross-entropy (log-loss). Uncertainty is quantified using 60% confidence intervals obtained via 500 bootstrap resamples of the test set.

For Fig. 4b, we evaluate generalization across the continuous genetic ancestry space. For each phenotype, we restrict in-context exemplars to individuals in the European (EUR) ancestry group (AoU provides discrete ancestry assignments for all subjects, the elastic net was trained from scratch on these eur samples) and evaluate predictions on test samples from the meta-ancestry cohort, ensuring that test individuals are, on average, further from the center of the training ancestry distribution (Ding et al., 2023). To quantify this shift, we compute the Mahalanobis distance (Mahalanobis, 1936) of each test individual from the training ancestry distribution based on the AoU-provided ancestry coordinates.

Let 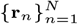 denote the ancestry-coordinate vectors of the training set. We compute the training-set mean

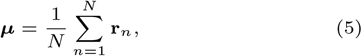

and a regularized inverse covariance matrix **Σ**^−1^ estimated from the training data. For a test individual with ancestry coordinate **r**, the Mahalanobis distance is

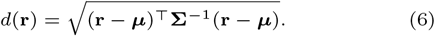

For both ICL-NS-G2P-proto and elastic net, we evaluate predictive performance as a function of ancestry distance by measuring, for each test subject, the log-probability assigned to the correct label. For Fig. 4b, to summarize the relationship between predictive performance and ancestry displacement, we partition test subjects into five uniformly sized bins based on their Mahalanobis distance from the center of the training ancestry distribution. The plot displays the mean (within a bin) log-probability of the correct label together with a 60% confidence interval of this statistic, computed via bootstrap resampling (500 bootstrap samples).

## Discussion

This study establishes that a central advantage of foundation models—their ability to operate effectively in sample-limited settings—extends to tabular foundation models applied to genomic risk prediction. Our results also identify a clear limitation of off-the-shelf tabular foundation models: they are not aligned to ancestry-dependent non-stationarity in genotype–phenotype relationships, a fundamental property of human genetic variation. This gap highlights substantial room for further alignment of tabular foundation models to the statistical realities of genomic data.

We refer to ICL-NS-G2P-proto as a prototype to emphasize that instruction tuning with synthetic data is only an initial step. Effective instruction tuning requires exposure to a sufficiently diverse family of tasks (Chan et al., 2022). Training exclusively on Gaussian process–based synthetic tasks over extended schedules risks over-adaptation to inductive biases specific to Gaussian Processes, rather than learning general strategies for handling non-stationary genetic architectures. Numerous alternative generative frameworks are well established in statistical genetics and bioinformatics for modeling ancestry-dependent variation in effect sizes, allele frequencies, sample availability, and linkage disequilibrium (Ma and Zhou, 2021; Moreno-Grau et al., 2024). Incorporating such task families during instruction tuning offers a path toward richer task diversity, longer training regimes, and improved alignment with real-world genotype–phenotype relationships.

## Funding & Acknowledgments

This work was supported by the US National Cancer Institute grant R01CA262296. We gratefully acknowledge All of Us participants for their contributions, without whom this research would not have been possible. We also thank the National Institutes of Health’s All of Us Research Program for making available the participant and sample data examined in this study. This study used data from the AoU Research Program’s Controlled Tier Dataset ver7, and ver8, available to authorized users. Ancestry coordinates and group labels used in this study are provided by the AoU Research Program for research purposes and reflect genetic principal component– based groupings; they should not be interpreted as proxies for race, ethnicity, or social identity. We emphasize the importance of responsible interpretation of ancestry-related findings in genomic research and caution against conflating genetic ancestry with social constructs of race.

## References

Y. Bai, C. Fan, H. Wang, C. Xiong, and S. Mei. Transformers as statisticians: Provable in-context learning with in-context algorithm selection. In Advances in Neural Information Processing Systems, 2023.

S. A. Bien, G. L. Wojcik, C. J. Hodonsky, C. R. Gignoux, I. Cheng, T. C. Matise, U. Peters, E. E. Kenny, and K. E. North. The future of genomic studies must be globally representative: Perspectives from page. Annual Review of Genomics and Human Genetics, 2019.

T. Brown, B. Mann, N. Ryder, M. Subbiah, J. D. Kaplan, P. Dhariwal, A. Neelakantan, et al. Language models are few-shot learners. Advances in Neural Information Processing Systems, 33:1877–1901, 2020.

S. Chan, A. Santoro, A. Lampinen, J. Wang, A. Singh, P. Richemond, J. McClelland, F. Hill, et al. Data distributional properties drive emergent in-context learning in transformers. Advances in Neural Information Processing Systems, 35:18878–18891, 2022.

S. H. Choi, X. Wang, E. A. Rosenthal, A. L. Blegen, S. J. Wirkus, V. A. Wagner, J. G. Meyer, M. S. Cicek, All of Us Research Program, et al. Genomic data in the all of us research program. Nature, 627(8003):340–346, 2024. doi: 10.1038/s41586-023-06957-x.

S. W. Choi, T. S. Mak, and P. F. O’Reilly. Tutorial: a guide to performing polygenic risk score analyses. Nature Protocols, 15:2759–2772, 2020. doi: 10.1038/s41596-020-0353-1.

Y. Cui. Digital pathways connecting social and biological factors to health outcomes and equity. npj Digital Medicine, 8:172, 2025. doi: 10.1038/s41746-025-01564-8.

A. Das, M. I. Khalid, R. Peñaloza, and S. Schockaert. When no paths lead to rome: Benchmarking systematic neural relational reasoning. Advances in Neural Information Processing Systems, 39, 2025.

Y. Ding, K. Hou, Z. Xu, A. Pimplaskar, E. Petter, K. Boulier, F. Privé, B. J. Vilhjálmsson, L. M. Olde Loohuis, and B. Pasaniuc. Polygenic scoring accuracy varies across the genetic ancestry continuum. Nature, 618:774–781, 2023. doi: 10.1038/s41586-023-06079-4.

L. Duncan, H. Shen, B. Gelaye, J. Meijsen, K. Ressler, M. Feldman, R. Peterson, and B. Domingue. Analysis of polygenic risk score usage and performance in diverse human populations. Nature Communications, 10:3328, 2019. doi: 10.1038/s41467-019-11112-0.

Y. Gao and Y. Cui. Deep transfer learning for reducing health care disparities arising from biomedical data inequality. Nature Communications, 11:1–8, 2020. doi: 10.1038/s41467-020-18918-3.

Y. Gao and Y. Cui. Optimizing clinico-genomic disease prediction across ancestries: a machine learning strategy with pareto improvement. Genome Medicine, 16:76, 2024. doi: 10.1186/s13073-024-01345-0.

Y. Gao, T. Sharma, and Y. Cui. Addressing the challenge of biomedical data inequality: An artificial intelligence perspective. Annual Review of Biomedical Data Science, 6:153–171, 2023.

S. Guerrero, A. López-Cortés, A. Indacochea, J. M. García-Cárdenas, A. K. Zambrano, A. Cabrera-Andrade, P. Guevara-Ramírez, D. A. González, P. E. Leone, and C. Paz-y Miño. Analysis of racial/ethnic representation in select basic and applied cancer research studies. Scientific Reports, 8:13978, 2018. doi: 10.1038/s41598-018-32264-x.

D. Gurdasani, I. Barroso, E. Zeggini, and M. S. Sandhu. Genomics of disease risk in globally diverse populations. Nature Reviews Genetics, 20:520–535, 2019. doi: 10.1038/s41576-019-0144-0.

A. Henry, X. Mo, C. Finan, M. D. Chaffin, D. Speed, H. Issa, S. Denaxas, et al. Genome-wide association study meta-analysis provides insights into the etiology of heart failure and its subtypes. Nature Genetics, 57:815–828, 2025. doi: 10.1038/s41588-024-02064-3.

N. Hollmann, S. Müller, L. Purucker, A. Krishnakumar, M. Körfer, S. B. Hoo, R. T. Schirrmeister, and F. Hutter. Accurate predictions on small data with a tabular foundation model. Nature, 637:319–326, 2025. doi: 10.1038/s41586-024-08328-6.

L. Kachuri, N. Chatterjee, J. Hirbo, D. J. Schaid, I. Martin, I. J. Kullo, E. E. Kenny, B. Pasaniuc, J. S. Witte, and T. Ge. Principles and methods for transferring polygenic risk scores across global populations. Nature Reviews Genetics, 2023. doi: 10.1038/s41576-023-00637-2.

I. J. Kullo. Clinical use of polygenic risk scores: current status, barriers and future directions. Nature Reviews Genetics, 2025. doi: 10.1038/s41576-025-00900-8.

D.-S. M. Lee, K. M. Cardone, D. Y. Zhang, N. L. Tsao, S. Abramowitz, P. Sharma, J. S. DePaolo, K. G. Aragam, S. B. Biddinger, et al. Common-variant and rare-variant genetic architecture of heart failure across the allele-frequency spectrum. Nature Genetics, 57:829–838, 2025. doi: 10.1038/s41588-025-02140-2.

N. J. Lennon, C. L. Abo, R. L. Chisholm, W. K. Chung, R. C. Green, G. P. Jarvik, I. J. Kullo, et al. Lessons learned from the emerge network: balancing genomics in discovery and practice. American Journal of Human Genetics, 109(6): 973–981, 2022. doi: 10.1016/j.ajhg.2022.05.002.

Y. Ma and X. Zhou. Genetic prediction of complex traits with polygenic scores: a statistical review. Trends in Genetics, 37:995–1011, 2021. doi: 10.1016/j.tig.2021.06.004.

P. C. Mahalanobis. On the generalised distance in statistics. Proceedings of the National Institute of Sciences of India, 2(1):49–55, 1936.

M. C. Mills and C. Rahal. The gwas diversity monitor tracks diversity by disease in real time. Nature Genetics, 52:242– 243, 2020. doi: 10.1038/s41588-020-0580-y.

S. Moreno-Grau, M. Vernekar, A. Lopez-Pineda, D. Mas-Montserrat, M. Barrabes, C. D. Quinto-Cortes, B. Moatamed, M. T. Lee, Z. Yu, K. Numakura, et al. Polygenic risk score portability for common diseases across genetically diverse populations. Human Genomics, 18:93, 2024. doi: 10.1186/s40246-024-00664-y.

R. E. Peterson, K. Kuchenbaecker, R. K. Walters, C.-Y. Chen, A. B. Popejoy, S. Periyasamy, M. Lam, C. Iyegbe, R. J. Strawbridge, L. Brick, et al. Genome-wide association studies in ancestrally diverse populations. Cell, 179:589–603, 2019. doi: 10.1016/j.cell.2019.08.051.

F. Privé, H. Aschard, S. Carmi, L. Folkersen, C. Hoggart, P. F. O’Reilly, and B. J. Vilhjálmsson. Portability of 245 polygenic scores when derived from the uk biobank and applied to 9 ancestry groups from the same cohort. American Journal of Human Genetics, 109:12–23, 2022. doi: 10.1016/j.ajhg.2021.11.008.

T. Sharma, N. K. Verma, and Y. Cui. Omics-based cancer prognosis across ethnic groups: From feature engineering to disparity detection and mitigation. IEEE Transactions on Artificial Intelligence, 2026. In press.

N. Shrine, A. G. Izquierdo, J. Chen, R. Packer, R. J. Hall, A. L. Guyatt, C. Batini, R. J. Thompson, C. Pavuluri, V. Malik, et al. Multi-ancestry genome-wide association analyses improve resolution of genes and pathways influencing lung function and chronic obstructive pulmonary disease risk. Nature Genetics, 55:410–422, 2023. doi: 10.1038/s41588-023-01314-0.

G. Sirugo, S. M. Williams, and S. A. Tishkoff. The missing diversity in human genetic studies. Cell, 177:26–31, 2019. doi: 10.1016/j.cell.2019.02.048.

C. Tcheandjieu, X. Zhu, A. T. Hilliard, S. L. Clarke, V. Napolioni, S. Ma, K. M. Lee, H. Fang, F. Chen, Y. Lu, et al. Large-scale genome-wide association study of coronary artery disease in genetically diverse populations. Nature Medicine, 28:1679–1692, 2022. doi: 10.1038/s41591-022-01891-3.

Y. Wang, J. Guo, G. Ni, J. Yang, P. M. Visscher, and L. Yengo. Theoretical and empirical quantification of the accuracy of polygenic scores in ancestry divergent populations. Nature Communications, 11:3865, 2020. doi: 10.1038/s41467-020-17719-y.

Y. Wang, M. Kanai, T. Tan, M. Kamariza, K. Tsuo, K. Yuan, W. Zhou, Y. Okada, BioBank Japan Project, H. Huang, et al. Polygenic prediction across populations is influenced by ancestry, genetic architecture, and methodology. Cell Genomics, 3:100408, 2023. doi: 10.1016/j.xgen.2023.100408.

J. Wei, M. Bosma, V. Y. Zhao, K. Guu, A. W. Yu, B. Lester, N. Du, A. M. Dai, and Q. V. Le. Finetuned language models are zero-shot learners. In International Conference on Learning Representations, 2022.

W. Zhou, M. Kanai, K.-H. H. Wu, H. Rasheed, K. Tsuo, J. B. Hirbo, Y. Wang, A. Bhattacharya, H. Zhao, S. Namba, et al. Global biobank meta-analysis initiative: Powering genetic discovery across human disease. Cell Genomics, 2:100192, 2022. doi: 10.1016/j.xgen.2022.100192.

