## Supplementary Information for "Bridging Ancestry Gaps in Genomic Risk Prediction with Tabular Foundation Models"

*Supplementary Information*, 2026, pp. 1–4

doi: Not assigned
Supplementary Information

SUPPLEMENTARY INFORMATION

### Supplementary Information for Bridging Ancestry Gaps in Genomic Risk Prediction with Tabular Foundation Models

Anirban Das[1,*] and Yan Cui[2,*]

[1]Department of Genetics, Genomics & Informatics, The University of Tennessee Health Science Center, Memphis, 38163, Tennessee, USA and [2]Department of Genetics, Genomics & Informatics, The University of Tennessee Health Science Center, Memphis, 38163, Tennessee, USA

#### Abstract

This document provides supplementary material for the paper titled *Bridging Ancestry Gaps in Genomic Risk Prediction with Tabular Foundation Models*.

#### Supplementary Content

#### Continuous ancestry coordinates

The All of Us (AoU) Research Program provides each participant with a 16-dimensional genetic principal-component (PC) coordinate computed from genome-wide genotype data (Choi et al., 2024). These coordinates are the same representation AoU uses internally to assign broad ancestry group labels (e.g., European, African, East Asian). Because the grouping is derived from thresholds applied to these very coordinates, using the full 16-dimensional vector as a continuous covariate retains strictly more information than the discrete labels: any function of the discrete label can be recovered from the coordinates, but not vice versa. In particular, within-group variation along ancestry PCs—which discrete labels discard—can capture fine-scale population structure that influences allele frequencies and, consequently, genotype–phenotype relationships (Ding et al., 2023).

In our pipeline, the 16-dimensional PC vector is concatenated with the genetic dosage features for each individual, so that predictive models can exploit the full ancestry continuum rather than collapsing it into a small number of categories. For the eMERGE dataset, where AoU-provided coordinates are not available, we compute principal components from genome-wide genotype data using PLINK and attach a 16-dimensional coordinate vector following the same convention.

#### Details of phenotype, target ancestry, and source ancestry triplets studied in Fig. 1 of the main paper

Summarized in Fig S1

#### eMERGE data wrangling pipeline

For the eMERGE III dataset, prior GWAS summary statistics are not available (Lennon et al., 2024; Zouk et al., 2019). We therefore perform phenotype-specific GWAS analyses using PLINK (Purcell et al., 2007), following a standardized multi-step pipeline applied exclusively to the training data to prevent data leakage. GWAS and linkage disequilibrium (LD) pruning are performed in a pooled manner using samples from all ancestry groups. Fifteen percent of samples are held out as an independent test set.

Step 1: Phenotype and covariate construction.
Case–control phenotypes are defined using curated eMERGE III clinical files. Disease status is mapped from dbGaP encodings (Mailman et al., 2007), and demographic data are merged to extract sex and birth year. Where applicable, a single body mass index (BMI) measurement is selected per subject. This step produces PLINK-compliant phenotype, covariate, and cohort inclusion files.

Step 2: Genotype subsetting and conversion.
Genotypes for selected subjects and chromosomes are extracted from imputed VCF files and converted to PLINK2 PGEN format, with explicit sample filtering applied during conversion.

Step 3: GWAS (training set only).
Association testing is performed using logistic regression with Firth correction (Wang, 2014) under an additive genetic model, adjusting for covariates. This step uses only genetic data and phenotype labels from the training set and produces full GWAS summary statistics.

Step 4: Statistical filtering.
Variants are filtered based on statistical significance ($P < 10^{-4}$) and allele frequency constraints ($0.01 \leq$ minor allele frequency $\leq 0.99$), yielding a reduced set of candidate variants.

| | phenoname | description | reference_ancestry | target_ancestry | heritability_target | heritability_reference |
|---|---|---|---|---|---|---|
| 0 | CA_120.1 | Myeloid | eur | afr | 0.000 | 0.079 |
| 1 | CA_120.1 | Myeloid | eur | amr | 0.049 | 0.079 |
| 2 | CA_138 | Benign neoplasm of the skin | eur | afr | 0.050 | 0.024 |
| 3 | CA_138 | Benign neoplasm of the skin | eur | amr | 0.034 | 0.024 |
| 4 | CA_138 | Benign neoplasm of the skin | eur | eas | 0.012 | 0.024 |
| 5 | CA_139.5 | Lipoma | eur | afr | 0.029 | 0.052 |
| 6 | CA_139.5 | Lipoma | eur | amr | 0.014 | 0.052 |
| 7 | CV_411 | Other diseases of pericardium | eur | afr | 0.029 | 0.021 |
| 8 | CV_411 | Other diseases of pericardium | eur | amr | 0.067 | 0.021 |
| 9 | CV_416.11 | Supraventricular tachycardia | eur | afr | 0.064 | 0.000 |
| 10 | CV_416.11 | Supraventricular tachycardia | eur | amr | 0.084 | 0.000 |

**Fig. S1.** The All-by-All study released by the All of Us (AoU) Research Program performs genome-wide association studies (GWAS) separately for every phenotype–ancestry pairing. AoU provides discrete ancestry labels for participants with whole-genome sequencing data. The All-by-All study reports the heritability of each phenotype within each ancestry group, which constitutes the primary source of the heritability estimates used in this work.

Step 5: Linkage disequilibrium pruning.

To remove redundant signals, variants are LD-pruned using a 250 kb window and an $r^2$ threshold of 0.1. Only training set samples are used for LD pruning, resulting in a biologically independent feature set.

Step 6: Dosage extraction and dataset assembly.

Additive dosages for the pruned variants are extracted and merged with covariates and phenotype labels to form the final machine learning–ready dataset. Only the top 50 most significant variants, ranked by GWAS $p$-value, are used to construct the tabular dataset.

Finally, principal components are computed using PLINK, and a 16-dimensional ancestry coordinate is attached to each sample to represent ancestry as a continuous space, consistent with the All of Us representation. Once the training and test tabular datasets are complete, elastic net and TabPFN models are constructed and evaluated using the same procedures as for the All of Us data.

#### Validation of the AoU data wrangling pipeline

The output of the AoU data wrangling pipeline (Fig. S3) is a collection of tabular training and test datasets constructed for each selected phenotype–ancestry pair. Each dataset contains genetic dosage features, continuous ancestry coordinates, phenotype labels, and standard covariates, including age and sex. To ensure that downstream model performance reflects genetic signal rather than covariate-driven effects, we additionally construct matched datasets in which cases and controls are balanced with respect to covariates and covariates are excluded from the input feature set.

For each phenotype–ancestry dataset, we train two supervised baseline models and fine-tune an off-the-shelf TabPFN model on the training data, followed by evaluation on held-out All of Us (AoU) version 8 test subjects. This procedure is repeated across six distinct phenotype–ancestry combinations. As shown in Fig. S2, all models achieve predictive performance largely above chance when using genetic dosage features alone on the test set, validating the integrity of the multi-stage data curation and feature construction pipeline. Moreover, relative model performance across phenotypes on version 8 test subjects is consistent with ancestry-specific heritability estimates reported for AoU version 7.

#### Gaussian process–based synthetic data generation for non-stationary genotype–phenotype modeling

To analyze the behavior of in-context learning models under ancestry-dependent non-stationarity, we generate synthetic genotype–phenotype datasets inspired by large-scale biobank studies. The data generation framework follows principles used in population-scale association studies, while explicitly introducing controlled non-stationarity in allele effect sizes across a continuous ancestry space.

##### Feature representation

Each synthetic individual is represented by three components:

1. **Ancestry coordinates.** Each individual is assigned a continuous ancestry coordinate

$$\mathbf{r} \in \mathbb{R}^A,$$

where $A$ denotes the dimensionality of ancestry space (e.g., principal components summarizing genome-wide variation).

2. **Genetic dosage features.** Each individual has a vector of additive genotype dosages

$$\mathbf{x} = (\eta_1, \eta_2, \dots, \eta_N) \in \mathbb{R}^N,$$

where $\eta_i \in \{0, 1, 2\}$ denotes the dosage of the $i$-th single-nucleotide polymorphism (SNP).

3. **Phenotype label.** A binary phenotype $Y \in \{0, 1\}$ is generated and observed only for in-context exemplars.

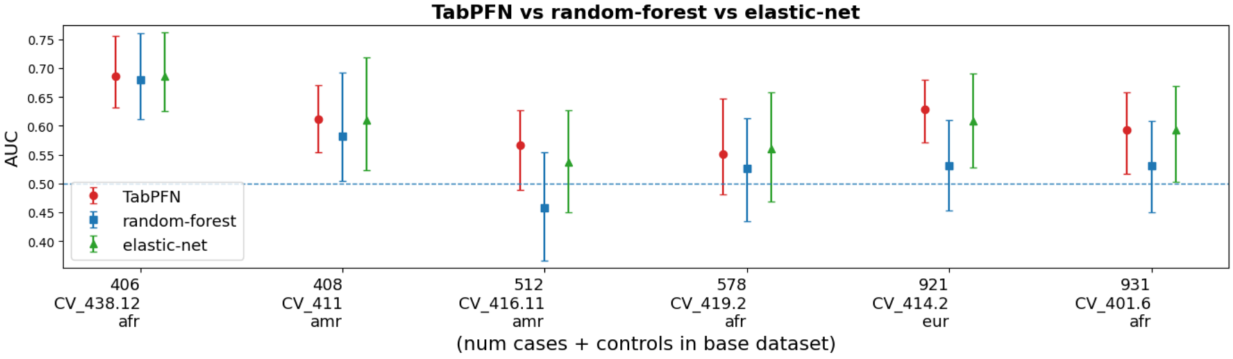


**Fig. S2.** Performance of TabPFN, Random Forest, and Elastic Net models for predicting multiple cardiovascular phenotypes across stratified ancestry groups. The horizontal blue line denotes the 0.5 baseline; confidence intervals entirely above this line indicate greater than 80% probability that the model captures predictive signal from genetic features alone. TabPFN models were fine-tuned for approximately 20 epochs (less than one minute). Elastic Net and Random Forest models were trained in a supervised manner and evaluated on held-out data. The AoU phenotype codes are CV_438.12 (Thoracic aneurysm), CV_411 (Other diseases of pericardium), CV_416.11 (Supraventricular tachycardia), CV_419.2 (Presence of cardiac defibrillator), CV_414.2 (Dilated cardiomyopathy), and CV_401.6 (Secondary hypertension).

##### Phenotype generation

Phenotypes are generated using a logistic regression model with ancestry-dependent effect sizes:

$$Y \sim \text{Bernoulli}\left(\sigma\left(\sum_{i=1}^{N} \beta_i(\mathbf{r})\,\eta_i\right)\right), \tag{1}$$

where $\sigma(\cdot)$ is the logistic sigmoid function. The functions $\beta_i(\mathbf{r})$ represent allele effect sizes that may vary across ancestry space and are not observed by the learning model.

##### Modeling non-stationary effect sizes

For a subset of $N_{\text{ctrl}}$ SNPs, effect sizes vary non-stationarily across ancestry space. These effect sizes are modeled as Gaussian processes:

$$\beta_i(\mathbf{r}) \sim \mathcal{GP}(0, k(\mathbf{r}, \mathbf{r}')), \tag{2}$$

with a radial basis function (RBF) kernel

$$k(\mathbf{r}_1, \mathbf{r}_2) = \exp\left(-\frac{\|\mathbf{r}_1 - \mathbf{r}_2\|_2^2}{2\ell^2}\right), \tag{3}$$

where $\ell$ is the kernel length scale.

The remaining SNPs have stationary effect sizes that do not depend on $\mathbf{r}$. Increasing $N_{\text{ctrl}}$ introduces more loci with ancestry-dependent effects, while smaller values of $\ell$ induce sharper local heterogeneity across ancestry space, creating more challenging forms of non-stationarity.

#### Hierarchical Gaussian process model and task generation

To generate more biologically realistic synthetic tasks, we extend the Gaussian process framework using a hierarchical structure that captures ancestry stratification, allele frequency variation, and sample density imbalance.

##### Ancestry clusters

We model ancestry space $\mathbb{R}^A$ as consisting of $K$ clusters with centers

$$\mathbf{c}_1, \ldots, \mathbf{c}_K \in \mathbb{R}^A,$$

such that inter-cluster distances are of order $L > 0$. Each individual $n$ is assigned a cluster label $z_n \in \{1, \ldots, K\}$ and an ancestry coordinate

$$\mathbf{r}_n \mid z_n = k \sim \mathcal{N}\left(\mathbf{c}_k,\ (\varepsilon L)^2 \mathbf{I}\right), \tag{4}$$

where $\varepsilon \in (0, 1)$ controls within-cluster dispersion.

Cluster-specific sampling probabilities control relative sample densities across ancestry regions.

##### Baseline allele frequency structure

For each SNP $i$ and cluster $k$, a cluster-specific minor allele frequency

$$\theta_{i,k} \in (0, 1)$$

is specified. Conditional on cluster membership, genotype dosages are sampled as

$$\eta_{n,i} \mid z_n = k \sim \text{Binomial}(2, \theta_{i,k}). \tag{5}$$

This construction induces systematic allele frequency differences across ancestry clusters, while retaining individual-level variability.

##### Hierarchical non-stationary effect sizes

Effect sizes are generated as continuous functions over ancestry space using the Gaussian process model in Eq. (3). Even within a single cluster, effect sizes may vary smoothly with $\mathbf{r}$, allowing for within-cluster heterogeneity. Variants with shorter length scales exhibit substantial local variation, while large length scales approximate stationary effects.

Clusters thus represent regions of shared sampling density and baseline allele frequency structure, without imposing piecewise-constant genetic effects.

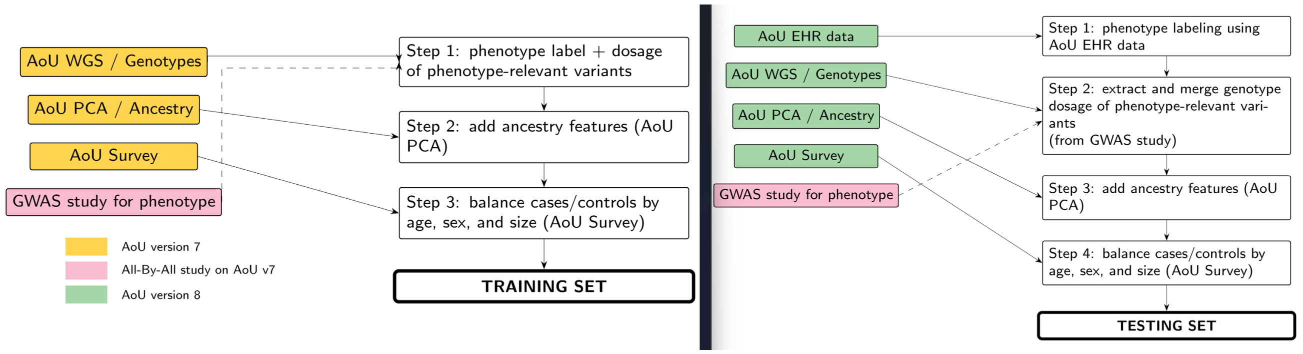


**Fig. S3.** Workflow illustrating how multiple primary data sources are integrated to produce feature–phenotype training and testing datasets suitable for testing TabPFN and related tabular foundation models. Each colored box is a different primary data source.

##### Task definition

Each unique configuration of hierarchical Gaussian process parameters

$$(K, A, L, \varepsilon, N_{\mathrm{ctrl}}, \ell)$$

defines a distinct synthetic genotype–phenotype prediction task. For each task, labeled and unlabeled samples are generated according to Eq. (1), and organized into tabular in-context learning problems consisting of labeled exemplars followed by unlabeled queries.

By varying these parameters, we generate a diverse family of tasks spanning different degrees of ancestry stratification, non-stationarity, and sample imbalance. These tasks form the basis of the analyses corresponding to Fig. 3 (left). For the analysis presented here, we set $\theta_{i,k} = \theta_i$ for all clusters $k$, such that all SNPs share the same call-rate across ancestry groups and no ancestry-dependent allele frequency differences are introduced.

#### Data curation pipeline for the All of Us dataset

Figure S3 summarizes the data curation workflow and the primary AoU resources linked by our pipeline. For each phenotype, the pipeline produces a tabular dataset whose rows correspond to participants and whose columns include: (1) allele dosages for a phenotype-specific set of variants, (2) continuous ancestry coordinates derived from AoU-provided 16-dimensional genetic principal components, (3) standard covariates such as age and sex, and (4) disease labels. The 16-dimensional principal component coordinates place each individual in a continuous genetic ancestry space and correspond to the same representation AoU uses internally to derive broad ancestry groupings, enabling fine-grained analyses of performance as ancestry varies continuously.

To prevent confounding due to differences in covariate distributions, we enforce matching of case and control groups within each dataset with respect to the marginal distributions of age and sex. In addition, the number of cases and controls is matched within both the training and test sets (Step 3 in Fig. S3). This design ensures that predictive performance reflects information extracted from genetic variation rather than from imbalanced covariate distributions.

#### References

S. H. Choi, X. Wang, E. A. Rosenthal, A. L. Blegen, S. J. Wirkus, V. A. Wagner, J. G. Meyer, M. S. Cicek, All of Us Research Program, et al. Genomic data in the all of us research program. *Nature*, 627(8003):340–346, 2024. doi: 10.1038/s41586-023-06957-x.

Y. Ding, K. Hou, Z. Xu, A. Pimplaskar, E. Petter, K. Boulier, F. Privé, B. J. Vilhjálmsson, L. M. Olde Loohuis, and B. Pasaniuc. Polygenic scoring accuracy varies across the genetic ancestry continuum. *Nature*, 618:774–781, 2023. doi: 10.1038/s41586-023-06079-4.

N. J. Lennon, L. C. Kottyan, C. Kachulis, N. S. Abul-Husn, J. Arias, G. Belbin, and J. E. Below. Selection, optimization and validation of ten chronic disease polygenic risk scores for clinical implementation in diverse us populations. *Nature Medicine*, 30, 2024.

M. D. Mailman, M. Feolo, Y. Jin, M. Kimura, K. Tryka, R. Bagoutdinov, L. Hao, et al. The ncbi dbgap database of genotypes and phenotypes. *Nature Genetics*, 39, 2007.

S. Purcell, B. Neale, K. Todd-Brown, L. Thomas, M. A. Ferreira, D. Bender, J. Maller, et al. Plink: a tool set for whole-genome association and population-based linkage analyses. *American Journal of Human Genetics*, 81, 2007.

X. Wang. Firth logistic regression for rare variant association tests. *Frontiers in Genetics*, 5, 2014.

H. Zouk, E. Venner, N. J. Lennon, D. M. Muzny, D. Abrams, S. Adunyah, L. Albertson-Junkans, et al. Harmonizing clinical sequencing and interpretation for the emerge iii network. *American Journal of Human Genetics*, 2019.
